# The mGluR5 Agonist, CHPG, Enhances Numbers of Differentiated Human Oligodendrocytes

**DOI:** 10.1101/2025.05.31.656838

**Authors:** Yangyang Huang, Celine Geywitz, Anjalika Bandaru, Ian A. Glass, Birth Defects Research Laboratory, Lucas Schirmer, Hiroko Nobuta, Cheryl Dreyfus

**Affiliations:** Department of Neuroscience and Cell Biology, Rutgers University, Piscataway, NJ 08854, USA; School of Environmental and Biological Sciences, Rutgers University, Piscataway, NJ 08854, USA; Center for Advanced Biotechnology and Medicine, Rutgers University, Piscataway, NJ 08854, USA; Interdisciplinary Center for Neurosciences, Heidelberg University, Mannheim, Germany; Division of Neuroimmunology, Department of Neurology, Heidelberg University, Mannheim, Germany; Medical Faculty Mannheim, Mannheim Center for Translational Neuroscience, Heidelberg University, Mannheim, Germany; University of Washington, Seattle, WA 98105, USA

**Keywords:** Oligodendrocyte, multiple sclerosis, demyelination, remyelination, induced pluripotent stem cells, postmortem tissues, human primary cells

## Abstract

Previous studies on adult mice indicate that the mGluR5 agonist 2-chloro-5-hydroxyphenyl glycine (CHPG), reduces cuprizone-elicited losses in myelin. This effect is partly mediated by CHPG binding to mGluR5 receptors on reactive astrocytes, triggering the release of brain derived neurotrophic factor (BDNF), which results in an increase in myelin, and alleviates behavioral deficits. However, it remains unclear whether CHPG has similar beneficial effects on human cells. To address this issue, we examined effects of CHPG on human cells using both human induced pluripotent stem cell (hiPSC)-derived oligodendrocytes and primary human fetal brain cells. Treatment of hiPSCs (30μM, 5 days) or primary cells (30 μM, 3 days) with CHPG increases the percent of MBP^+^O4^+^ mature oligodendrocytes relative to total O4^+^cells, without affecting survival. When effects of CHPG were evaluated on proliferating OPCs, effects on proliferation are observed. In contrast, when CHPG was evaluated in young oligodendrocytes, effects on proliferation were gone, suggesting that in this population CHPG is influencing differentiation. Interestingly, in contrast to observations in mice, mGluR5 expression in humans is localized on PDGFRα^+^ oligodendrocyte precursor cells (OPCs) and O4^+^ immature oligodendrocytes, but not astrocytes. Moreover, using purified human OPC cultures, we show a direct effect of CHPG in enhanced differentiation. To identify potential cellular targets of CHPG in the <u>adult</u> human brain, we analyzed postmortem tissue from individuals with multiple sclerosis (MS) and healthy controls. In contrast to the hiPSCs or fetal cells, demyelinated white matter from MS patients showed elevated mGluR5 mRNA expression in astrocytes. Taken together, our findings suggest that CHPG enhances the differentiation of human OPCs during development through a mechanism distinct from that observed in adult cuprizone-treated mice. Moreover, astrocytes in MS pathology upregulate mGluR5, suggesting they may become responsive to CHPG under disease conditions.

## Introduction

In people with multiple sclerosis (MS), restoration of lost myelin is considered a key step toward functional recovery. Pathological features of MS suggest some evidence of remyelination following myelin breakdown [1, 2], but it may be incomplete or inefficient [1, 3]. Oligodendrocyte precursor cells (OPCs) are abundant in both acute and chronic lesions as well as periplaque white matter (PPWM) [4–7], suggesting that a block in differentiation rather than a lack of progenitor cells may underlie remyelination failure. These observations suggest that facilitating differentiation in OPCs may be a plausible strategy for restoring myelination in MS.

We have previously shown in mice that the group I/II metabotropic glutamate receptor (mGluR) agonist trans-(1S,3R)-1-amino-1,3-cyclopentanedicarboxylic acid (ACPD) induced the release of BDNF from astrocytes in culture [8] and *in vivo* following a cuprizone lesion [9]. BDNF binds to TrkB receptors on oligodendrocyte lineage cells, which in turn differentiate and increase myelin protein production and myelination [10, 11]. Subsequently, we showed using the highly specific mGluR5 agonist CHPG and antagonist 2-methyl-6-(phenylethynyl) pyridine (MPEP) that these effects are mediated by mGluR5 signaling [10]. Moreover, the source of BDNF was found to be astrocytes in cuprizone-fed mice, because astrocyte-specific deletion of BDNF or astrocyte-specific deletion of mGluR5 was sufficient to reverse the effects of CHPG-mediated effects on oligodendrocyte lineage cells [10]. Interestingly, control-fed mice did not respond to CHPG, suggesting cuprizone-induced reactive astrocytes may be required [10].

The current study investigated whether the beneficial role of CHPG on oligodendrocytes can be observed in human oligodendrocyte lineage cells using two human cellular models (human induced pluripotent stem cell (hiPSC)-derived oligodendrocytes and primary human fetal brain cell culture). We provide evidence that CHPG induces a significant increase in the proportion of mature oligodendrocytes and the number of primary processes emanating from the oligodendrocyte cell body. These effects were reversed by the specific mGluR5 antagonist MPEP. In contrast to adult lesioned mice, mGluR5 expression in developing human cells was restricted to neurons and OPCs and not in astrocytes. Isolated human OPCs responded to CHPG and increased the proportion of mature oligodendrocytes, demonstrating a direct effect of CHPG on OPC differentiation. In contrast to the developing human cellular models, adult postmortem tissues from multiple sclerosis (MS) and healthy controls revealed that whereas mGluR5 expression in astrocytes in healthy white matter was low, a significant upregulation was observed in MS lesions, implying a phenotypic change in astrocytes in the presence of a lesion, consistent with upregulation of mGluR5 in cuprizone treated adult mice. Together these observations indicate that, while species differences may exist in the expression patterns of mGluR5, CHPG in a developing system may directly mediate OPC differentiation while in a lesioned adult the effects of CHPG are through astrocytes.

## Results

### CHPG enhances numbers of differentiated human oligodendrocytes

We generated human OPCs from hiPSCs using a previously established protocol [12] that results in the production of neurons and astrocytes in addition to oligodendrocytes (Fig. 1A). Cultures were treated with the mGluR5 agonist CHPG (30μM) for 5 days, either from culture day 35 to 40, when only young oligodendrocyte precursors are present (Fig. 1B), or from day 50 to 55, when O4^+^ immature oligodendrocytes are present (Fig. 1C) [12]. Regardless of treatment timing, all analyses were performed at day 55, a stage at which both MBP^+^ mature oligodendrocytes and MBP^-^/O4^+^ immature oligodendrocytes are present. Because of our previous data indicating that BDNF enhances levels of MBP in mouse cultured oligodendrocyte lineage cells [13], we also defined effects of this neurotrophin on the human cells.

**Figure 1.**
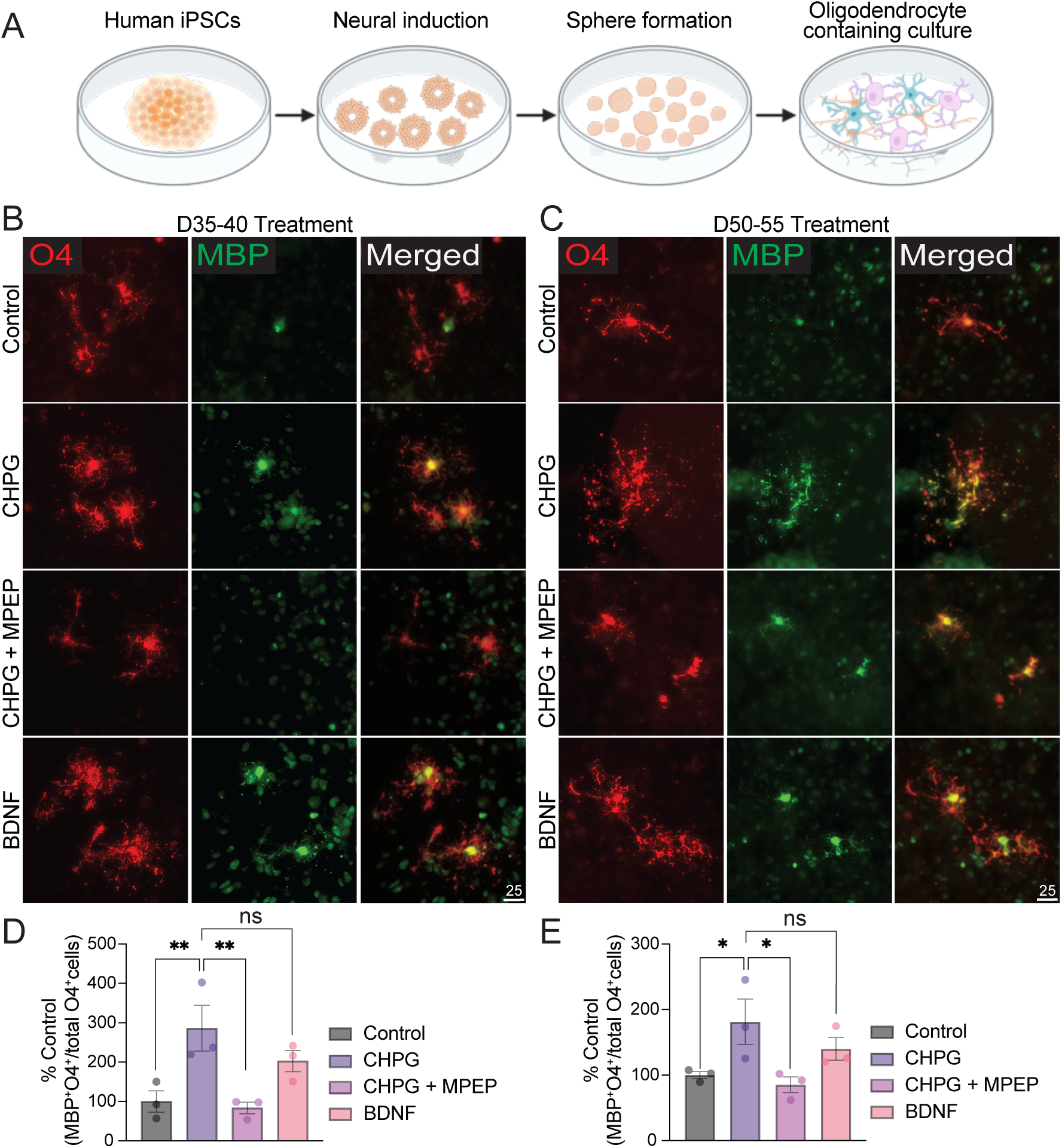
CHPG enhances numbers of differentiated human oligodendrocytes derived from iPSCs. (**A**) Schematic illustration of human iPSC-derived oligodendrocyte culture. (**B-C**) CHPG (30μM final concentration) alone, or CHPG together with mGluR5 antagonist MPEP (20μM final concentration), or BDNF (10ng/ml final concentration) was added during pre-OL stage (day 35-40, panel **B**) or immature-to-mature OL stage (day 50-55, panel **C**). (**D-E**) The proportion of mature OL, defined by the percentage of MBP^+^O4^+^ double positive cells relative to all O4^+^ cells, was quantified on day 55, when cultures contain mature OLs. CHPG significantly increased the proportion of mature OL at both time points, but its effect was bigger during day 35-40 (N = 3 independent experiments). Both effects were reversed by MPEP. Effects of BDNF did not differ from those of CHPG. Scale bar = 25μm.

When the hiPSC-derived culture were treated with CHPG, we observed a significant increase in the proportion of MBP^+^O4^+^ mature oligodendrocytes relative to total O4^+^ cells, reaching levels comparable to those induced by BDNF (10ng/ml) (Fig. 1B-E). Although the enhanced expression of MBP was observed regardless of the timing of CHPG application, day 35 to 40 showed higher significance compared to day 50 to 55 (p = .0061, D35-40 vs p = .0232, D50-55, Fig. 1D-E). These effects were reversed by mGluR5 antagonist MPEP (20μM), indicating mGluR5 dependent action of CHPG (Fig. 1D-E).

To further validate our findings, we used an additional human model of mixed glial cultures derived from fetal brain tissue, which contain neurons, astrocytes, and oligodendrocytes (Suppl Fig. 1, Fig. 2A), and assessed the effects of CHPG on oligodendrocyte differentiation. As indicated in Figure 2B-D, and consistent with results from iPSC-derived cultures (Fig. 1), a 3-day treatment with CHPG enhanced the proportion of mature oligodendrocytes and promoted the number of primary processes emanating from the oligodendrocyte cell body. These effects were reversed by co-treatment with MPEP (Fig 2B-D).

**Figure 2.**
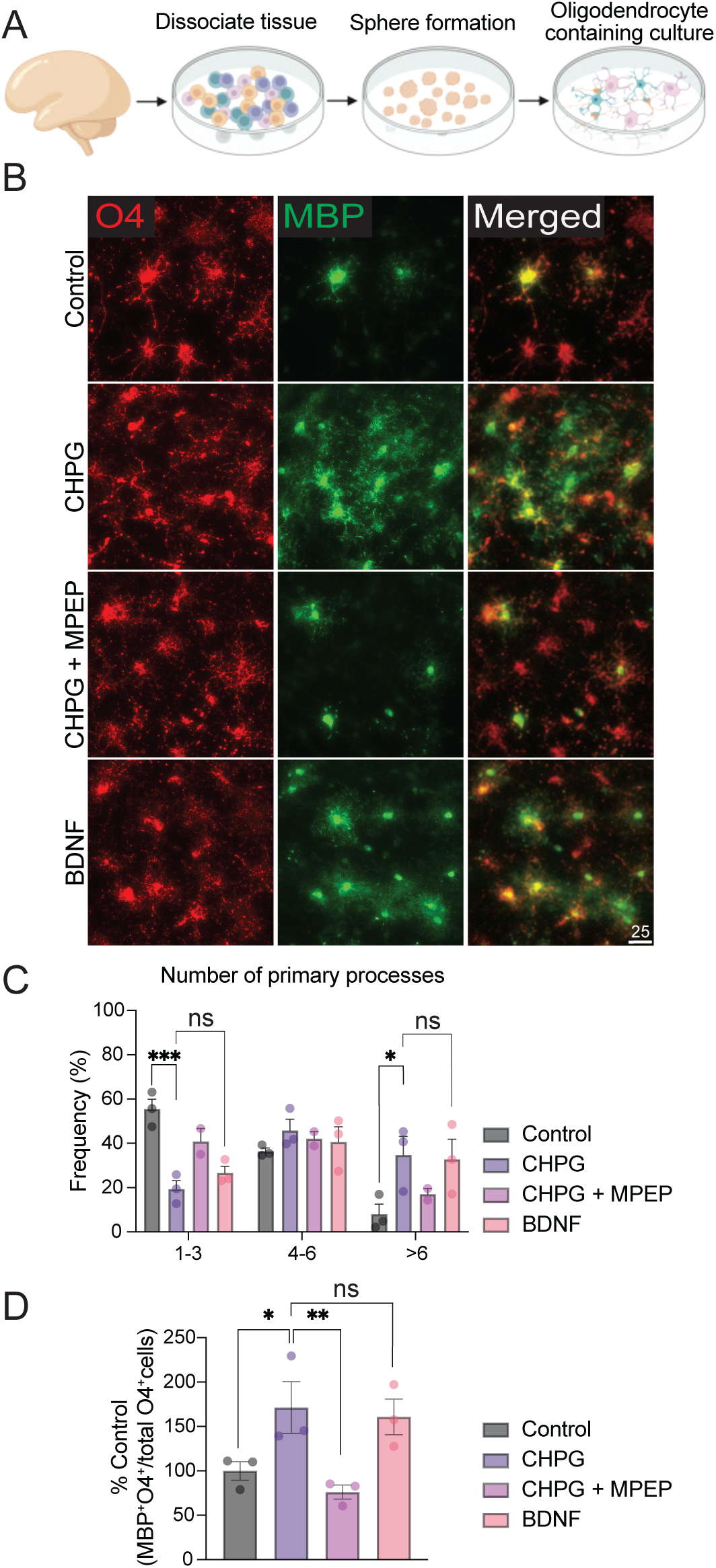
CHPG enhances numbers of differentiated human primary oligodendrocytes. (**A**) Schematic illustration of the derivation of human mixed glial culture from primary brain tissues. (**B**) Human brain cell culture containing neurons, astrocytes, and OPCs were treated with CHPG (30μM final concentration) alone, or CHPG together with mGluR5 antagonist MPEP (20μM final concentration), or with BDNF (10ng/ml final concentration) for 3 days in Glia medium (see Methods). (**C**) The number of primary processes was counted for every O4^+^ cell and expressed as percent of total cells exhibiting 1-2, 4-6 or >6 processes. (**D**) The proportion of mature OL, defined by the percentage of MBP^+^O4^+^ double positive cells relative to all O4^+^ cells, was quantified. CHPG treatment significantly increased the proportion of mature OL, which was reversed by MPEP (N = 3 independent experiments). Effects of BDNF did not differ from those of CHPG. Scale bar = 25μm.

### CHPG increases OPC proliferation in a developmentally regulated manner but has no effect on survival

The previous findings that CHPG enhanced numbers of mature oligodendrocytes at both early (day 35-40) and later (day 50-55) stages of development (Fig. 1B-E) suggests that CHPG may influence proliferation, survival or differentiation. To assess whether CHPG affects OPC proliferation, we added EdU, a marker of DNA synthesis, during CHPG treatment at both the early pre-OL (day 35-40) and later immature-to-mature OL stage (day 50-55) of hiPSC-derived differentiating oligodendrocytes. On day 55, we quantified Edu^+^O4^+^ oligodendrocytes from both time windows. Results indicate that CHPG treatment during the early (day 35-40) stage significantly increased numbers of EdU^+^O4^+^ oligodendrocytes (Suppl Fig. 2A, C), whereas CHPG treatment during the later (day 50-55) stage did not affect the numbers of EdU^+^O4^+^ cells (Suppl Fig. 2B, D). Overall, these findings suggest that CHPG affects proliferation in young, but not older cultured cells’. Moreover, the survival of oligodendrocytes was not affected at any OPC developmental stage by CHPG, as detected by caspase-3 immunofluorescent staining (Suppl Fig. 3A-B) or propidium iodide inclusion (data not shown).

### mGluR5 is expressed in human neurons and immature oligodendrocytes but not in astrocytes or mature oligodendrocytes

To investigate the cell types that are responsive to CHPG, we used immunofluorescent staining for mGluR5 on human primary cultures. Consistent with previous mouse studies [14–17], we detected mGluR5 expression in MAP2^+^ and TUJ1^+^ neurons (Fig. 3A) as well as PDGFRα^+^ OPCs and O4^+^ immature oligodendrocytes (Fig. 3B), while MBP^+^ mature oligodendrocytes were negative for mGluR5 (Fig. 3B). Interestingly, GFAP^+^ human astrocytes lacked mGluR5 expression (Fig. 3A). Publicly available gene expression data on human brain cells [18–20] as well as Human Protein Atlas proteinatlas.org confirmed minimal expression of mGluR5 (gene name *GRM5*) in human astrocytes under healthy conditions. These findings indicate that in unlesioned human cells, CHPG likely acts directly on OPCs and immature oligodendrocytes rather than astrocytes. These results suggest an additional mechanism of action of CHPG in unlesioned human cells to those noted in cuprizone-demyelinated adult mouse, in which the binding of CHPG to mGluR5 expressed on mouse astrocytes triggers the release of BDNF to facilitate OPC differentiation [9, 10].

**Figure 3.**
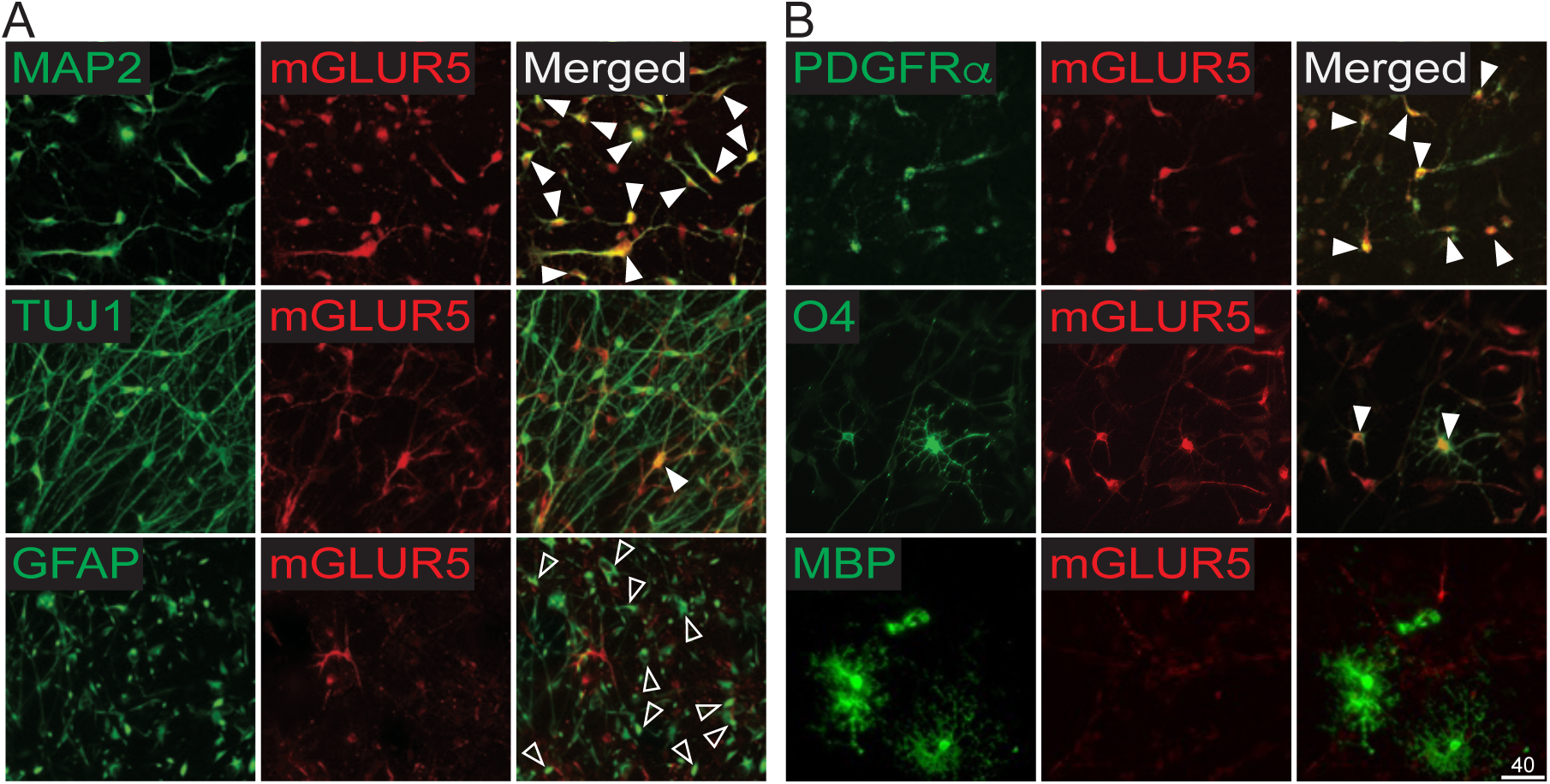
mGluR5 expression in human brain cells. (**A**) mGluR5 antibody and cell type specific antibodies were used to immunofluorescently determine mGluR5 expression. MAP2^+^ and TUJ1^+^ neurons express mGluR5 (solid arrowheads). GFAP^+^ astrocytes are negative for mGluR5 (empty arrowheads). (**B**) in the OL lineage cells, only PDGFRα^+^ OPC and O4^+^ immature cells express mGluR5 (solid arrowheads), but not MBP^+^ mature OLs. Scale bar = 40μm.

### CHPG directly enhances human OPC differentiation

The evidence that human immature oligodendrocytes express mGluR5 (Fig. 3B) suggests a direct effect of CHPG on these cells. Therefore, we isolated PDGFRα^+^ OPCs using immunopanning (see methods) from human primary mixed glia cultures (Fig 4A). The isolated cells were 92% pure (PDGFRα^+^) with minor contaminants from astrocytes. These cultures were then treated with CHPG for 5 days and the proportion of mature oligodendrocytes was quantified. The results indicate a significant increase in the proportion of MBP^+^O4^+^ mature oligodendrocytes in the CHPG-treated group, reaching levels similar to those observed with BDNF (Fig 4B-C). The effect was reversed by MPEP (Fig 4C), indicating that CHPG directly promotes the differentiation of human OPCs through mGluR5.

**Figure 4.**
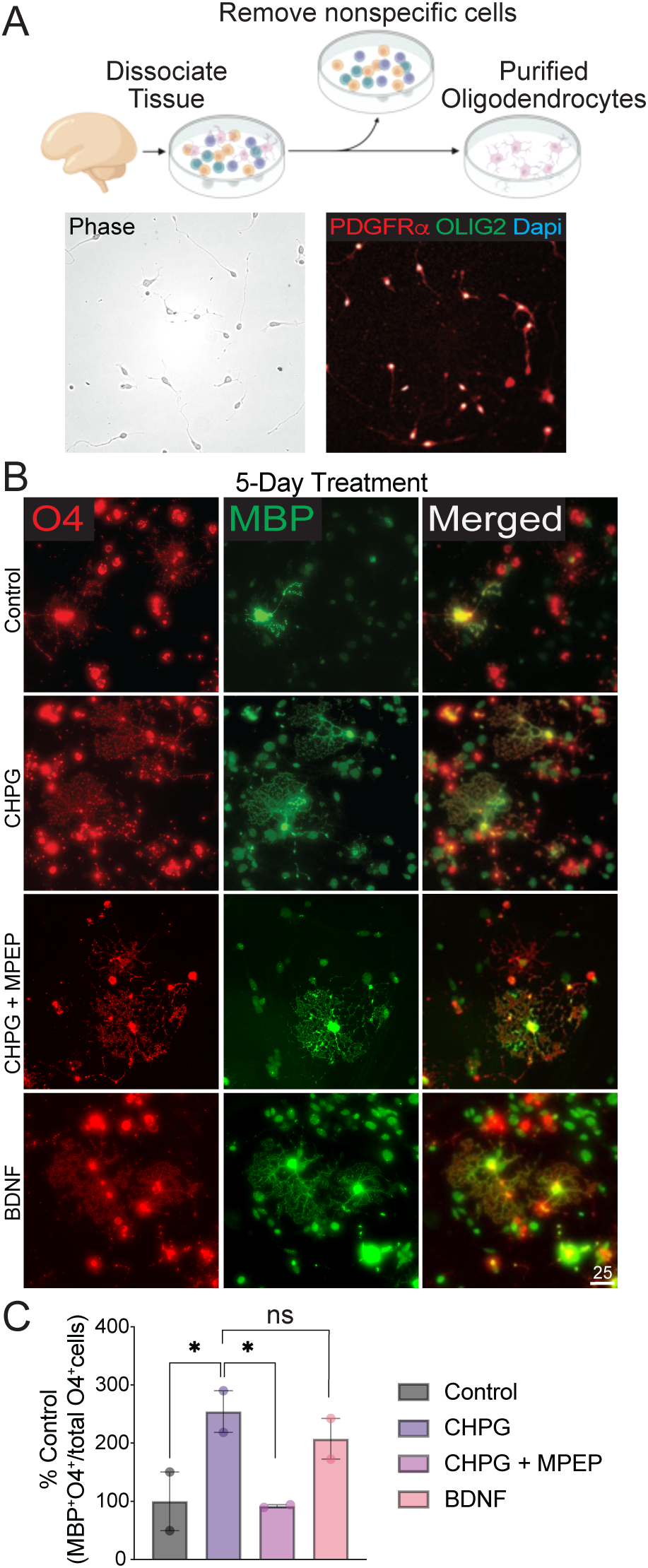
CHPG directly enhances human OPC differentiation. (**A**) Schematic illustration of PDGFRα^+^ human OPCs isolation by modified immunopanning. (**B**) Purified human OPCs were treated with CHPG (30μM final concentration) alone, or CHPG together with the mGluR5 antagonist MPEP (20μM final concentration), or with BDNF (10ng/ml final concentration) for 3 days in differentiation medium. (**C**) The proportion of mature OL, defined by the percentage of MBP^+^O4^+^ double positive cells relative to all O4^+^ cells, was quantified. CHPG treatment significantly increased the proportion of mature OL, which was reversed by MPEP (N = 2 independent experiments). Effects of BDNF were not significantly different from those of CHPG. Scale bar = 25μm.

#### Postmortem tissues from MS show increased mGluR5 expression in astrocytes and decreased expression in OPCs

Our observations on adult mice suggested beneficial effects of CHPG in promoting remyelination in demyelinating diseases. To determine if this might be the case in adult humans, we investigated mGluR5 expression in postmortem brain tissues from four MS patients and four healthy controls (Suppl. Table 1).

Within the MS tissue, we examined three distinct regions: normal-appearing white matter (NAWM), peri-plaque white matter (PPWM), and demyelinated white matter (DMWM), which were determined by MOG immunohistochemistry (IHC) (Fig. 5A). After lesion characterization, we performed in-situ hybridization (ISH) using probes for *GRM5* and *SOX9* to assess co-expression of mGluR5 in astrocytes. We successfully detected signals from all probes in all tissue samples tested (Fig. 5B-C, Fig. 6A). We then quantitated the percent of astrocytes expressing *GRM5* in all regions (Fig. 5D). As expected, we found minimal expression of *GRM5* in astrocytes in control brains as well as in NAWM of MS samples, consistent with Figure 3 and previous reports [18–20] (Fig. 5D). In contrast, our results revealed an increase in the percent of *GRM5^+^/SOX9^+^*cells in the PPWM and DMWM compared to the NAWM (Fig. 5D). Next, we assessed mGluR5 expression on *PCDH15*^+^ OPCs, within the same post-mortem brain samples from MS patients and healthy controls. Our ISH data revealed that the percent of OPCs expressing mGluR5 decreased in the DMWM compared to the NAWM and PPWM of the MS tissues (Fig. 5E). Taken together, while mGluR5 expression in astrocytes increased in the PPWM and DMWM compared to NAWM, its expression in OPCs was notably reduced in the DMWM, suggesting a cell-type specific shift in mGluR5 expression within MS lesions.

**Figure 5.**
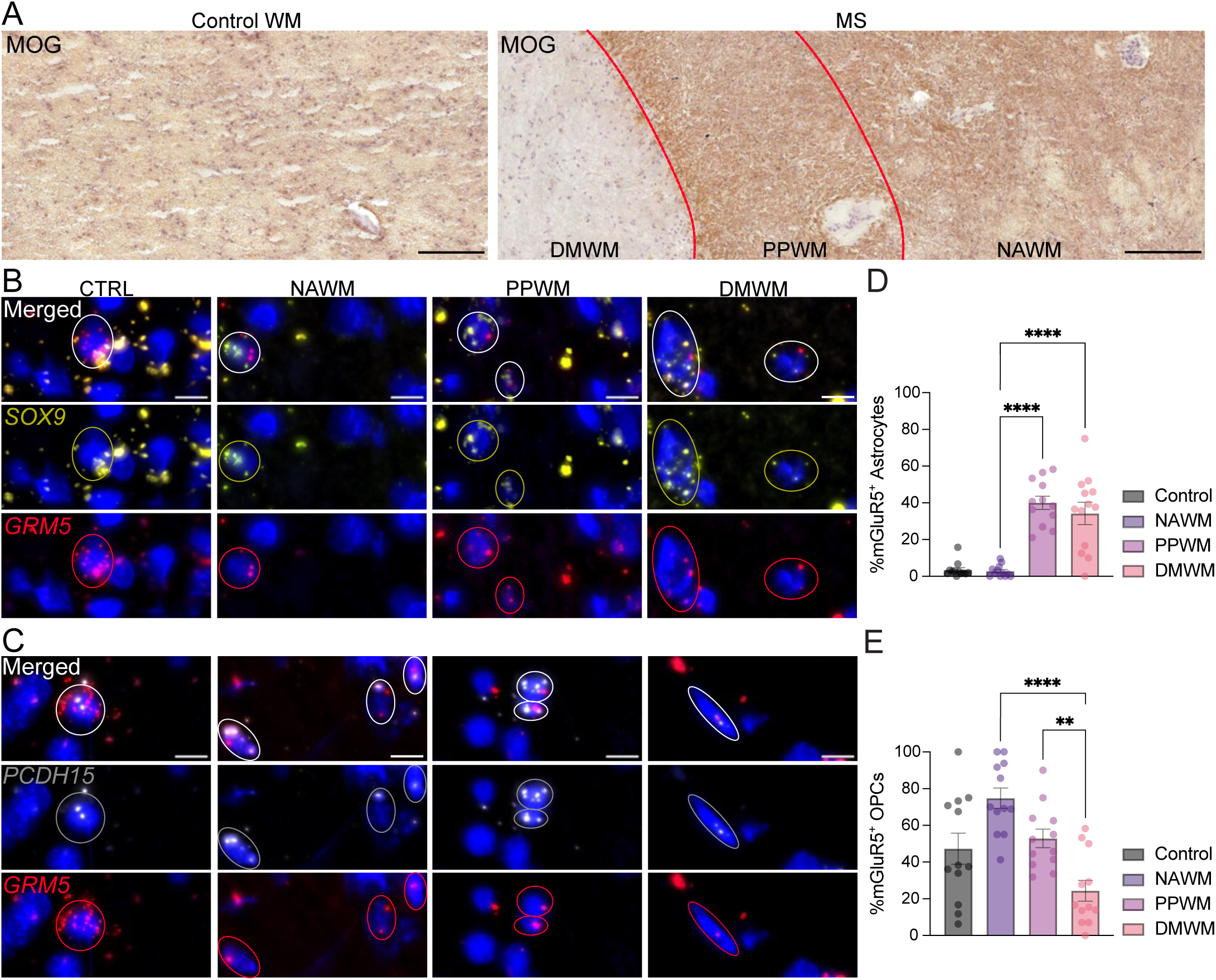
Differential expression of mGluR5 (*GRM5*) in astrocytes and OPCs in MS. (**A**) MOG immunohistochemistry was used to delineate borders between normal-appearing white matter (NAWM), peri-plaque white matter (PPWM), and demyelinated white matter (DMWM). *In situ* hybridization images of (**B**) *SOX9*^+^ (for astrocytes)/*GRM5*^+^ and (**C**) *PCDH15*^+^ (for OPCs)/*GRM5*^+^ cells in CTRL (N = 4) and chronic active MS lesions (N = 4). Circled dapi in **B, C** are examples of positive cells. (**D, E**) statistical analyses of percentage of double-positive cells from 6 regions of interest/sample. Scale bar = 10μm.

**Figure 6.**
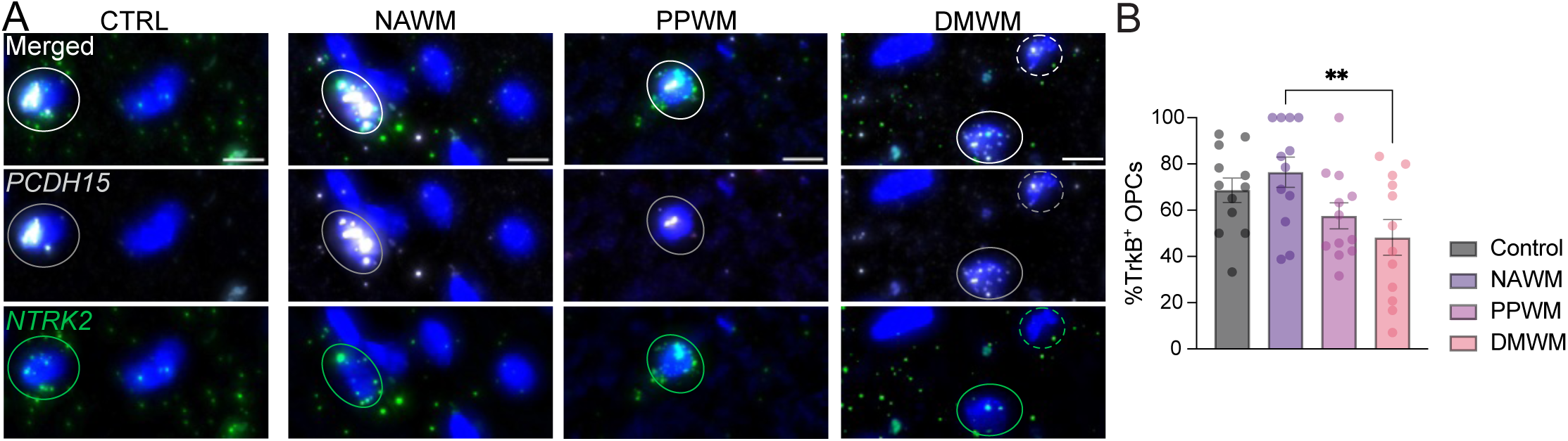
Differential expression pattern of TrkB (*NTRK2*) in OPCs in MS. (**A**) *In situ* hybridization images of *PCDH15*^+^ (for OPCs)/*NTRK2*^+^ cells in CTRL (N = 4) and chronic active MS lesions (N = 4). Circled dapi in **A** are examples of *NTRK2*^+^cells. Dotted dapi in **A** is an example of *NTRK2*^-^ cell. (**B**) Statistical analyses of percentage of double-positive cells from 6 regions of interest/sample. NAWM: normal-appearing white matter. PPWM: peri-plaque white matter. DMWM: demyelinated white matter. Scale bar = 10μm.

In cuprizone treated mice, we found that stimulation of astrocytes with CHPG increased levels of BDNF, which then interacted with TrkB receptors on oligodendrocyte lineage cells to enhance maturation [9, 10]. Accordingly, we assessed the expression of TrkB (*NTRK2*) in OPCs. Our findings revealed the presence of *NTRK2^+^/PCDH15^+^* cells in all three MS regions including NAWM, PPWM, and DMWM (Fig. 6A). Remarkedly, 50% of OPCs continue to express TrkB within the DMWM (Fig. 6B). These data suggest that despite *NTRK2* downregulation in the lesion center, many OPCs can respond to BDNF.

## Discussion

Previous studies have indicated that in a cuprizone-treated mouse the mGuR5 agonist CHPG elicits the synthesis and release of BDNF from astrocytes, resulting in the reversal of deficits in myelin proteins, myelin itself, and behavioral deficits. The deletion of mGluR5 from astrocytes, or the deletion of BDNF from astrocytes, or the deletion of trkB from oligodendrocyte lineage cells abrogates these effects [9–11], suggesting our working model that in an **adult mouse** subjected to a demyelinating lesion, CHPG stimulates BDNF from astrocytes to enhance oligodendrocyte recovery.

This study was designed to examine whether the observations noted in mice are relevant to human. We used two human models, hiPSC-derived oligodendrocytes [12], and human primary fetal cells (Suppl Fig. 1), both of which contain OPCs and mature oligodendrocytes, but recapitulate developmental stages rather than adult brain. Initial experiments examining the effects CHPG, targets of CHPG action, as well as the mechanism underlying the CHPG effects on oligodendrocyte differentiation revealed that CHPG does elicit differentiation of OPCs (Fig. 1B-E, Fig. 2B-D), but the effects are directly on the OPCs and not through astrocytes (Fig. 4B-C), which do not exhibit CHPG receptor mGluR5 in these models (Fig. 3A). However, in the adult MS brain, astrocytes within the lesion do exhibit mGluR5 and the proportion of mGluR5^+^ astrocytes appears to increase in comparison to healthy control or unlesioned MS brain (Fig. 5D-E). To our knowledge, these data are the first to demonstrate the regulation of mGluR5 expression at RNA (in situ) level with distinct periplaque and lesion mapping. It is unknown what effect stimulation of mGluR5^+^ astrocytes would have on adult OPCs in the lesion site or on myelination, but it is interesting that they express TrkB, suggesting a mechanism as seen in the adult mouse where stimulation of astrocytes to release BDNF in turn can stimulate remyelination, although other astrocyte-derived molecules may also be involved.

Since astrocytes in adult human MS lesions, as well as OPCs themselves express the CHPG receptor mGluR5 (Fig. 5D-E), the data suggest that the cells responsible for the effects of CHPG to influence OPC differentiation may differ as animals mature or are exposed to pathological influences. During development, OPCs directly respond to CHPG without astrocytic contribution. In adulthood, in the presence of cuprizone demyelinating pathologies, astrocytes mediate the effects of CHPG.

The literature supports this possibility that cells responding to CHPG change during maturation and in response to a lesion. mGluR5 protein and transcript are highly expressed throughout the rodent brain during development [16, 21] and decrease in adulthood [22, 23]. With respect to mGluR5 expression on oligodendrocyte lineage cells, a number of studies report that rodent [17] and human pre-myelinating oligodendrocytes express mGluR5 [24] only at specific times during their development in culture and *in vivo,* although this may vary in specific CNS regions [25]. Moreover, stimulation of these receptors affects oligodendrocyte lineage cell function. For example, an mGluR5 agonist, 3,5-dihydroxyphenylglycine (DHPG), reduces oligodendrocyte lineage cell death in culture [16, 21, 26] and the general Group I mGluR agonist ACPD, as well as DHPG, protects oligodendrocyte lineage cells from death *in vivo* from demyelinating pathologies [24, 25].

Expression of mGluR5 on astrocytes appears to be developmentally regulated as well. For example, it is reported in rodents that mGluR5 is highly expressed during development on astrocytes in the first two postnatal weeks of development but is lost as animals mature [27, 28]. The decrease in mGluR5 is correlated with a loss in responsiveness to DHPG [27]. Upon injury, however, astrocytic mGluR5 is upregulated in rodent brains, for example following epilepsy or seizure [29–31], or mechanical injury [32].

The expression of mGluR5 on human astrocytes has not been thoroughly explored, but it has been reported that, similar to the situation in rodents, mGluR5 mRNA is low in adult astrocytes from unlesioned brain [18, 19, 28]. However, in the present work we found a strong mGluR5 upregulation in MS lesion-associated astrocytes (Fig. 5D-E), similar to a study that reported GFAP^+^ astrocytes within human MS lesions upregulate mGluR5 protein as defined by immunohistochemistry [33]. Also, it has been reported that astrocytes proximate to Alzheimer’s disease amyloid beta plaques [34] and in spinal cords of amyotrophic lateral sclerosis [35] upregulate mGluR5. Our current studies are consistent with these observations and support the possibility that human astrocytes do not express mGluR5 in the mature, healthy brain, but expression is elevated in periplaque and demyelinated MS lesion areas.

When we evaluated TRKB, we confirmed a robust *NTRK2* (encoding the BDNF receptor TRKB) expression in human OPCs in control and MS NAWM. However, we found a gradual *NTRK2* downregulation in periplaque and MS lesion areas (Fig. 6B). While these findings indicate that BDNF signaling might be altered in MS lesion areas, it is important to note that 48% of OPCs in MS lesion still express *NTRK2* (Fig. 6B). The possibility therefore remains that BDNF may play a role in these *NTRK2*^+^ OPCs.

In summary, our studies suggest that mGluR5 agonists may influence human oligodendrocyte differentiation during the lifetime of individuals by employing two mechanisms of action. During development these agonists have direct actions on OPCs to enhance maturation. We suggest that mGluR5 agonists may directly impact OPCs to influence conditions such as white matter injury in preterm infants [36–38], or juvenile MS [39–41] where white matter deficits are evident. However, in adults, effects on lesioned CNS may be mediated through actions on astrocytes. The finding that mGluR5 is present on astrocytes in MS lesions (Fig. 5E) is consistent with our previous work in mice that CHPG can enhance myelination after a demyelinating lesion through this intermediary. It remains to be seen if CHPG analogs may also enhance remyelination in MS or other conditions of demyelination.

## Acknowledgement

This work was supported by the Congressionally Directed Medical Research Programs through the MSRP under Award No. W81XWH-22-1-0788, awarded to C.D. Opinions, interpretations, conclusions, and recommendations are those of the authors and are not necessarily endorsed by the Department of Defense. H.N. acknowledges Birth Defects Research Laboratory (BDRL) for human brain tissue collection. BDRL is funded by NIH under NICHD Grant # R24HD000836, to I.A.G. H. N. acknowledges the generous support from the Amy P. Goldman Foundation. Schematic illustrations included in Figures 1, 2, 4, and Supplementary Figure 1 were created using BioRender by H.N. (agreement numbers OI286GYT76, IY286H0PV6, KN286H0KMJ).

## Methods

### Human iPSC Lines

Human iPSC cell lines used in this study are line # 051121-01-MR-017 and # 051104-01-MR-040 distributed by New York Stem Cell Foundation via a material transfer agreement, and a wildtype line described in [1]. A normal karyotype was confirmed in all lines. Mycoplasma contamination has been confirmed negative in all iPSC lines.

### OPC differentiation from human iPSCs

A previously published protocol for directed iPSC differentiation to OPCs [2] was used with the following modifications: human ES cell medium and human ES cell medium without basic FGF were used in place of mTeSR and custom mTeSR, respectively. Human ES cell medium contained: Knockout DMEM (ThermoFisher), 20% Knockout serum replacement (ThermoFisher), 1X glutamax (ThermoFisher), 1X NEAA (ThermoFisher), FGF2 (Peprotec, final concentration 10ng/ml). The concentration of SAG was 0.5 μM, T3 was 40ng/ml, NT3 was 1ng/ml. Penicillin-Streptomycin was omitted from N2 medium, HGF from PDGF medium, and HEPES from Glia medium.

On day 0, iPSCs in passage 18 to 20 were plated at 0.125X10^6^/well in a matrigel-coated 6-well plate with ES medium without basic FGF, supplemented with dual SMAD inhibitors, RA, and ROCK inhibitor Y27632 (Reprocell). From day 1 to day 4, N2 medium was gradually increased by 25% each day, reaching 100% on day 4. On day 8, dual SMAD inhibitors were replaced with SAG. On day 12, cells were lifted, dissociated, and seeded on petri dishes for sphere formation. On day 20, the medium was changed to PDGF medium. On day 30, spheres were plated on poly-L-ornithine/laminin-coated dishes. On day 45, the medium was changed to Glia medium.

### Compound treatments

The following drugs were added to OPC differentiation cultures from day 35 to 40 or day 50 to 55: 2-chloro-5-hydroxyphenylglycine (CHPG) (Tocris Bioscience, final concentration 30 μM), 2-methyl-6-(phenylethynyl) pyridine (MPEP) (Tocris Bioscience, final concentration 20 μM) brain derived neurotrophic factor (BDNF; Peprotec, final concentration 10ng/mL). Fifty percent of the medium containing the compounds was changed every other day.

### Human mixed glial culture from fetal brain tissues

Specimens from fetal (gestational weeks 17 to 19) human brain tissues were obtained from the Birth Defects Research Laboratory at the University of Washington with ethics board approval and maternal written consent. This study was performed in accordance with ethical and legal guidelines of the Rutgers University’s institutional review board. Fetal brain tissues were dissociated with papain for 10min. Single cell suspension was transferred to petri dish for 7 to 10 days to allow formation of neurospheres in PDGF medium described above. Neurorspheres were collected and plated onto coverslips to adhere and allow cells to migrate out. After 4 to 5 days, compound treatments were initiated in Glia medium described above. Cells were fixed after 3-day treatment for immunofluorescent staining.

### Isolation of human primary OPCs

Fetal brain tissues were dissociated with papain for 10min. The single cell suspension was passed onto a series of culture dishes coated with mouse IgG secondary antibody to avoid non-specific binding, and PDGFRα, to isolate PDGFRα OPCs. At the end of PDGFRα binding, adherent cells were washed, and detached with Trypsin, and collected by centrifugation. Cells were plated at 25,000 cells/well in 24-well plate and treated with compounds for 5 days, fixed, and immunofluorescently stained.

### Immunofluorescent staining

After the cells were fixed on day 55, immunocytochemistry was done with antibodies to O4, MBP, and Caspase-3, followed by image acquisition using the Leica DMi8 Microsystems fluorescent microscope. O4 is expressed in immature to mature oligodendrocytes and is a cell surface marker. MBP is an oligodendrocyte-specific protein that is only expressed in the cytoplasm of mature OLs. Caspase-3 is a cell death marker that is expressed in the cytoplasm of the cell.

### EdU proliferation assay

EdU was added during day 35 to 40, or 50 to 55 of the oligodendrocyte differentiation protocol from human iPSCs. On day 55, oligodendrocytes that proliferated during the respective time frame were quantified by EdU detection following the manufacturer’s protocol and co-immunofluorescently stained with O4 antibody.

### Quantitative Analysis

Fluorescent images were captured by Leica DMi8 microscope equipped with a sCMOS camera. 20X images taken from 4 technical replicates in each experiment were manually analyzed for the proportion of MBP^+^O4^+^ mature oligodendrocytes relative to all O4^+^ oligodendrocytes in randomly selected fields. At least 300 O4^+^ cells were used for quantification in each treatment group in each experiment. Numbers of EdU^+^O4^+^ cells were counted in randomly selected fields of 0.39mm^2^ and expressed as percent of EdU^+^O4^+^ cell numbers in cells grown under control conditions. Numbers of Caspase-3^+^O4^+^ cells were counted in randomly selected fields of 0.39mm^2^ and expressed as percent of Caspase-3^+^O4^+^ cell numbers in cells grown under control conditions. The number of primary processes extending from O4^+^ cell numbers were counted in randomly selected fields of 0.39mm^2^. Frequency was determined as percent of cells exhibiting 1-3, 4-6 or >6 processes. All analyses were conducted blindly of treatment groups.

### Antibodies

Following primary antibodies were used for staining.

Olig2 (Millipore Sigma AB9610)

O4 (Mouse hybridoma)

MBP (AbD Serotec MCA409S)

PDGFRa (BD Bioscience 556001)

GFAP (ThermoFisher 13-0300)

TUJ1 (Biolegend 801213)

MAP2 (Sigma M9942)

Caspase-3 (Cell Signaling 9664S)

Click-iT™ EdU Cell Proliferation Kit (ThermoFisher C10337)

Secondary antibodies of appropriate species were purchased from Jackson ImmunoResearch and Fisher Scientific.

### Human postmortem tissue samples

Human post-mortem brain samples with chronic active MS and healthy controls were obtained from the UK Multiple Sclerosis Tissue Bank at Imperial College London and received ethical approval from the National Research Ethics Committee in the UK (08/MRE09/31). A total of 8 post-mortem snap-frozen human brain blocks (4 MS and 4 control) were examined by performing multiplex RNA-scope in-situ hybridization.

### Immunohistochemistry staining human post-mortem brain tissue

For the classification of demyelination in post-mortem human brain tissue, sections were subjected to immunohistochemistry staining using the diaminobenzidine (DAB) kit (SK-4105, vector laboratories) to detect myelin oligodendrocyte glycoprotein (MOG). In brief, tissue sections were fixed at room temperature (RT) in ice cold 100% methanol for 5 min, followed by a wash in PBS. Subsequently, the tissue sections were outlined using a hydrophobic Pap pen (VEC-H-400, Biozol) and blocked in 10% Goat serum (16210064, Thermo Fisher) at RT. Post blocking, the sections were incubated with the MOG primary antibody (MAB5680, RRID: AB_1587278, Merck Millipore) at 4°C overnight. The following day, the sections were washed and incubated with a biotinylated secondary IgG antibody for 2 hours at RT. After washes, the sections were incubated with avidin-biotin complex for 1 hour at RT, the sections were then counterstained with 50% hematoxylin for 38 seconds. They were then rinsed and dehydrated through a graded ethanol series. Before mounting, the sections were cleared in 100% xylene. Finally, the sections were mounted using Eukitt (Orsatec).

### Multiplex RNA scope in-situ hybridization

RNAscope in-situ hybridization (ISH) was performed on control and MS post-mortem human brain samples with probes for human *GRM5* (122224, ACD), *NTRK2* (402621, ACD), *PCDH15* (525881, ACD), and *SOX9* (404221, ACD), all obtained from Advanced Cell Diagnostics (ACD). The RNAscope ISH protocol was performed following the manufacturer’s instructions with minor modifications (ACD, RNAscope Multiplex Fluorescent Detection Reagent v2, 323110).

Probes were diluted as stated in the protocol. Samples were hybridized for 2 hours at 40°C and washed twice in 1x wash buffer. The signal amplification was carried out by sequential incubations with v2Amp1 (30 min), v2Amp2 (30 min) and v2Amp3 (15 min) at 40°C, each followed by two washes. Subsequently, sections were treated with v2-HRP-C1 for 15 min at 40°C and then washed twice with 1x wash buffer.

TSA-conjugated fluorophores were diluted 1:500 in TSA buffer and incubated for 30 min at 40°C followed by 2 washes and a subsequent incubation with a HRP blocker for 30 min at 40°C. The last steps were performed subsequently for v2-HRP-C2 and v2-HRP-C3. Finally, the sections were mounted using ProLong Gold Antifade mounting medium (P36930, Thermo Fisher).

### Microscopy and quantification

After completing the RNAscope ISH protocol, stained sections from post-mortem human brains were imaged using the Leica DM6 B Thunder microscope, equipped with a Leica K5C camera. Images were acquired at 40x magnification to ensure detailed visualization of the target probes. All images were captured as z-stacks according to the Nyquist criteria to ensure comprehensive signal capture within the tissue sections.

Prior to imaging, different regions of the lesion were selected based on myelin oligodendrocyte glycoprotein (MOG) immunohistochemistry staining to identify specific anatomical landmarks and pathological features. The following regions were investigated: normal white matter (WM) of controls (CTRL), normal appearing WM (NAWM) at least 7000 µm from any discernible demyelinated lesion rim, white matter surrounding lesions (peri-plaque WM, PPWM), and demyelinated lesional areas (demyelinated WM, DMWM).

Subsequently, images were loaded into the Fiji/ImageJ software for processing. Maximum intensity projections were created from the z-stacks for analysis. Within each region (CTRL, NAWM, PPWM, DMWM), six regions of interest (ROIs, area: 51.25 mm^2^) were selected randomly. To avoid bias, all fluorescent channels except DAPI were turned off during ROI selection. Subsequently, positive and double-positive cells within each ROI were then manually counted. From these six ROIs, two random ROIs were averaged, resulting in three values per region.

### Statistical analysis

Each *in vitro* experiment except the experiment described in Figure 4 (purified human OPCs) was repeated at least three times. 3-to-4 biological replicates were included in each experiment. The experiment described in Figure 4 was repeated twice only, due to the scarcity of human OPCs that can be obtained from donated tissues. For statistical analysis, two sided One-way ANOVA and t-test with normal distribution, were used as appropriate.

Statistical analysis and plot generation of the human brain tissue were performed using Python. Quantitative data were analyzed to compare the mean percentage of double-positive cells between control (n=4) and MS (n=4) samples. Statistical analysis was performed using parametric (one-way ANOVA) or nonparametric (Kruskal-Wallis) tests. The following significance levels were used: * p < 0.05, ** p < 0.01, *** p < 0.001, **** p < 0.0001, ns = not significant.

## Data Availability

The data that support the findings of this study are available from the corresponding author upon reasonable request.

---

**Supplemental Figure 1.**
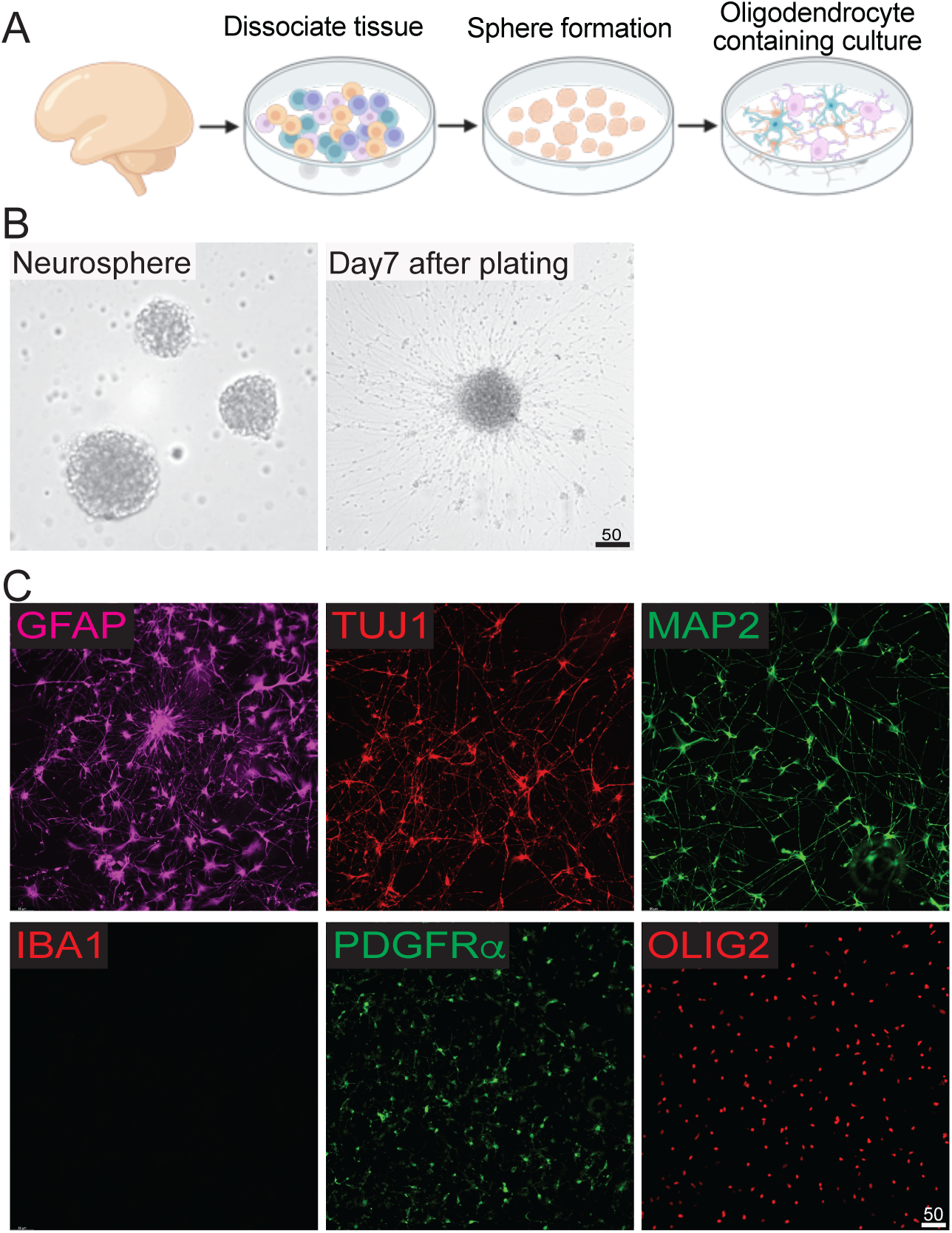
Human brain cell culture. (**A**) Fetal brain tissues (gestational weeks 17 to 19) were dissociated and allowed the formation of neurospheres in PDGF medium (see Methods) for 7-10 days, then plated to allow migration of cells for 4-5 days. (**B**) On the day of compound treatments, the culture contains GFAP^+^ astrocytes, TUJ1^+^, MAP2^+^ neurons, PDGFRα^+^, OLIG2^+^ OPCs. The culture is devoid of IBA1^+^ microglia. Scale bar = 50μm.

**Supplemental Figure 2.**
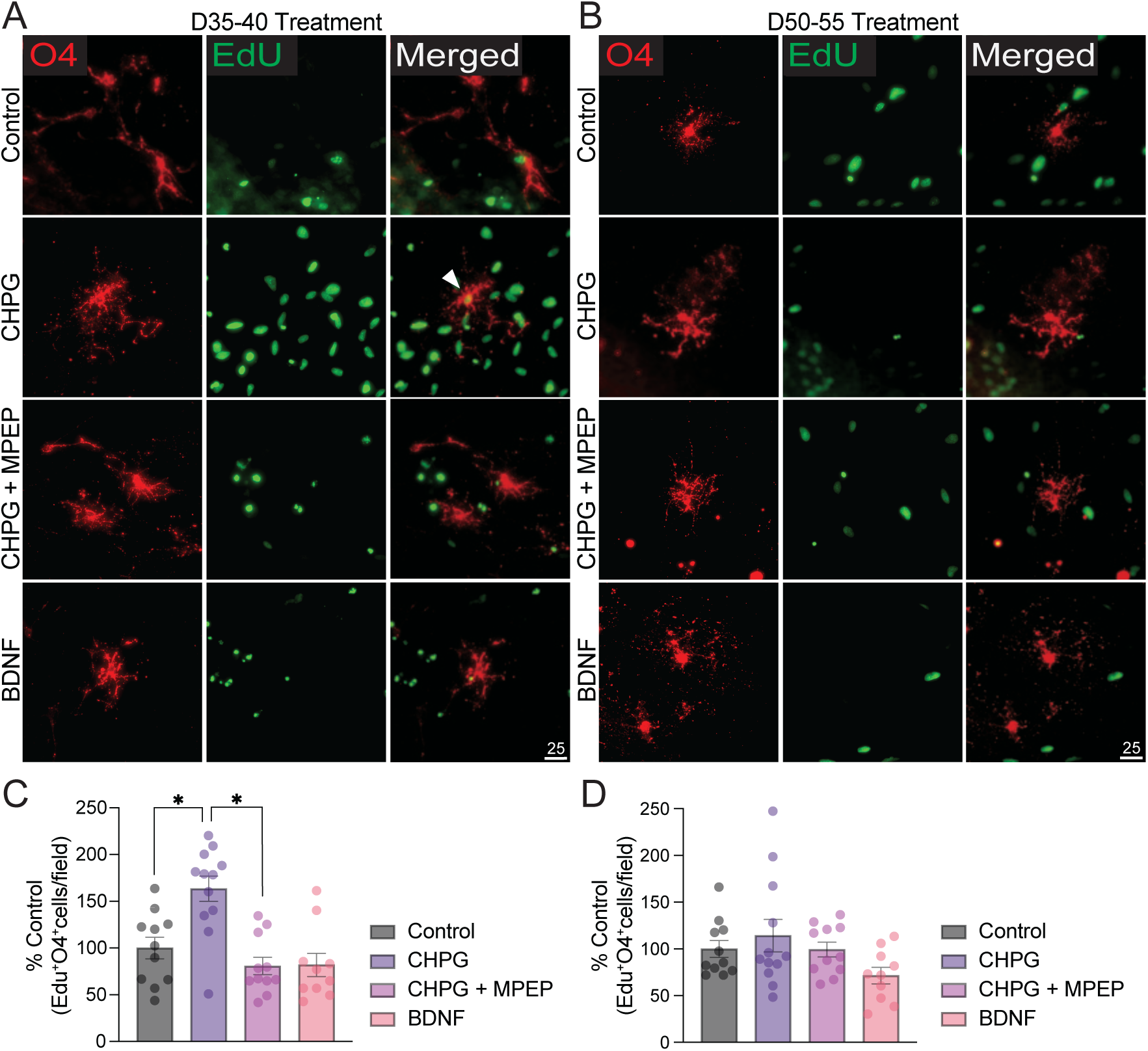
CHPG increases proliferation of pre-OLs. Edu was added during pre-OL stage (day 35-40, panel **A**) or immature-to-mature OL stage (day 50-55, panel **B**) of iPSC differentiation, concurrently with CHPG (30μM final concentration), CHPG+MPEP (20μM final concentration), or BDNF (10ng/ml final concentration) treatment. (**C-D**) On day 55, oligodendrocytes that proliferated were quantified by Edu detection and co-immunofluorescent staining with O4. CHPG significantly increased proliferation of OL lineage cells only when CHPG was given during pre-OL stage (day 35-40) (N = 3 independent experiments). The effect was reversed by MPEP. Scale bar = 25μm.

**Supplemental Figure 3.**
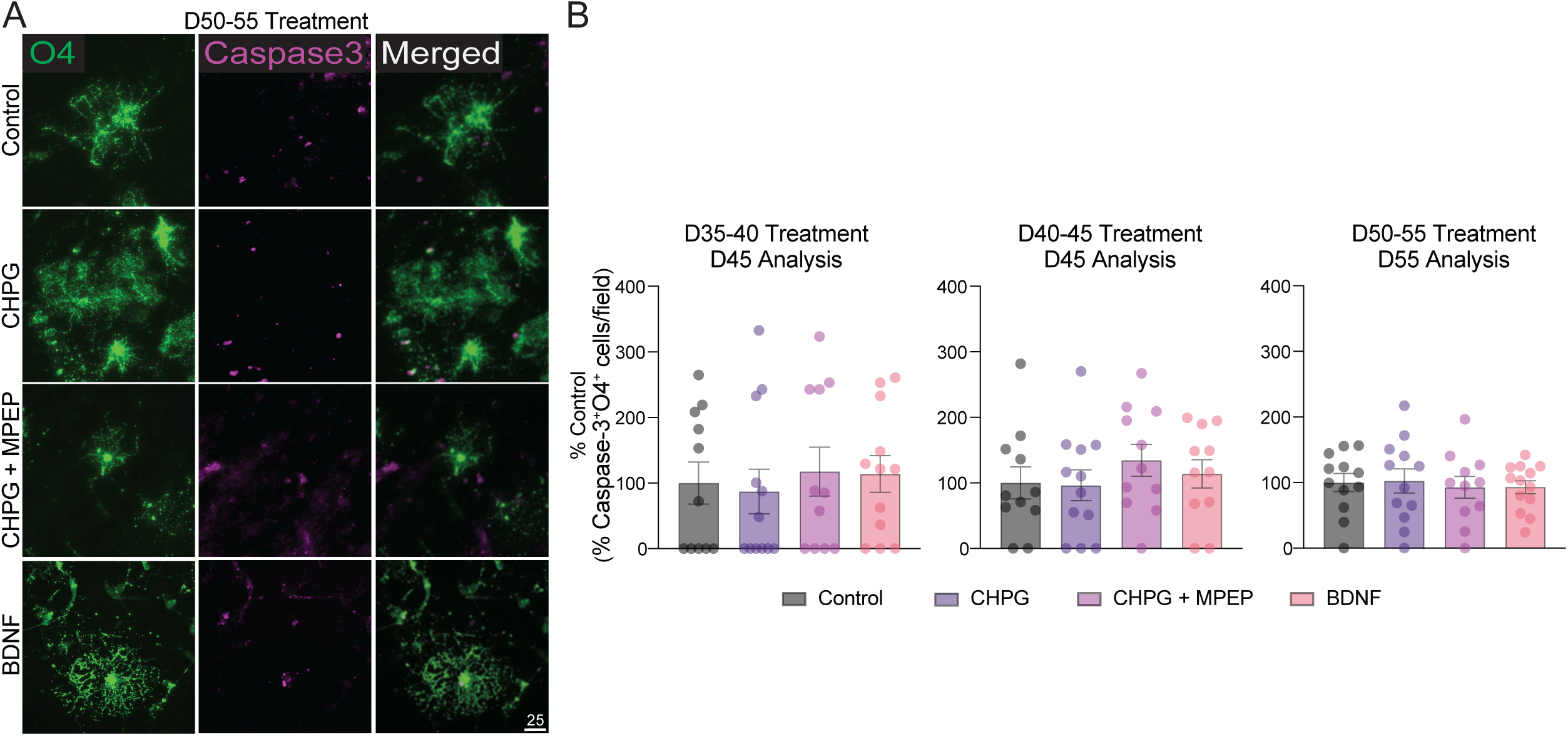
CHPG does not affect OL survival. (**A**) OL cell death was analyzed with Caspase-3 antibody at the end of CHPG (30μM final concentration) treatment at various stages of iPSC differentiation (day 35-40, 40-45, or 50-55) and colocalized with O4. (**B**) Overall cell death remained low throughout time points (<4% or all O4^+^ cells), and no significant difference was observed in response to CHPG, MPEP, or BDNF treatments (N = 3 independent experiments). Scale bar = 25 μm.

**Supplemental Table 1.** List of MS and control tissues used in this study.

| Sample ID | Age | MS status | Sex | RNA Integrity Number | Disease duration (years) | Lesion type |
| --- | --- | --- | --- | --- | --- | --- |
| CO22 | 69 | Control | female | 5.2 | NA | NA |
| CO64 | 63 | Control | female | 6 | NA | NA |
| CO90 | 83 | Control | male | 4.7 | NA | NA |
| PDCO40 | 61 | Control | female | 5.7 | NA | NA |
| MS426 | 48 | MS | female | 7 | 28 | chronic active |
| MS438 | 53 | MS | female | 5 | 18 | chronic active |
| MS513 | 51 | MS | male | 4.8 | 18 | chronic active |
| MS528 | 45 | MS | female | 5.2 | 25 | chronic active |

